# *In situ* targeted mutagenesis of gut bacteria

**DOI:** 10.1101/2022.09.30.509847

**Authors:** Andreas K Brödel, Loïc Charpenay, Matthieu Galtier, Fabien J Fuche, Rémi Terrasse, Chloé Poquet, Marion Arraou, Gautier Prevot, Dalila Spadoni, Edith M Hessel, Jesus Fernandez-Rodriguez, Xavier Duportet, David Bikard

## Abstract

Microbiome research is revealing a growing number of bacterial genes that impact our health. While CRISPR-derived tools have shown great success in editing disease-driving genes in human cells, we currently lack the tools to achieve comparable success for bacterial targets. Here we engineer a phage-derived particle to deliver a base editor and modify *E. coli* colonizing the mouse gut. This was achieved using a non-replicative DNA payload, preventing maintenance and dissemination of the payload, while allowing for an editing efficiency of up to 99.7% of the target bacterial population. The editing of a β-lactamase gene resulted in the stable maintenance of edited bacteria in the mouse gut at least 42 days after treatment. By enabling the *in situ* modification of bacteria directly in the gut, our approach offers a novel avenue to investigate the function of bacterial genes and provides an opportunity to develop novel microbiome-targeted therapies.

## Introduction

In recent years, microbiome research has unraveled an increasing number of mechanisms by which the expression of genes from commensal bacteria can impact our health. Bacteria can affect the success of immunotherapies^1–5^, bacterial antigens are involved as peptide mimics in autoimmune diseases^6,7^, bacterial toxins can drive a range of acute and chronic diseases including cancer^8,9^, bacteria can modify or sequester drugs impacting the effectiveness of therapies^10,11^. This growing list is driving interest in manipulating the microbiome, both to better understand it and to develop novel therapies.

Current strategies attempt to modify the gene repertoire by changing the microbiome composition through the use of broad or narrow antimicrobials, the addition of new strains, or through dietary changes^12^. These methods are confronted with the complexity of reliably and stably modifying microbial ecosystems of which we have a poor understanding. Here we propose a strategy to perform *in situ*, precise and stable genetic modifications of target bacterial populations rather than modifying the microbiome composition.

Achieving this goal in the complex environment of the gut requires an efficient DNA delivery strategy. Two main methods have been proposed so far to introduce DNA in bacteria of the gut microbiome: transduction by a bacteriophage capsid^13,14^, or conjugation from a donor bacterium^13,15,16^. Conjugation is attractive in that it can enable DNA delivery to a broad range of strains and species from a single donor strain. Transfer rates are however low for most recipients, and most conjugative systems work poorly in the gut environment^17^. High transfer efficiencies can be achieved in the animal gut, but rely on the stabilization of the mating pair through specific interactions between donor pili and receptors on the recipient surface^18^. Efficient conjugative delivery to different strains and species will thus likely require different specialized systems. Strategies relying on conjugation also suffer from the need to administer live genetically modified bacteria that will spread an engineered genetic circuit.

Bacteriophages have also been explored as delivery vehicles. Previous works on DNA delivery to *E. coli* in the mouse gut have relied either on M13 cosmids^14^, or on genetically modified bacteriophage λ^19,20^. M13 uses the F pilus as a receptor which limits its range, and M13 virions were shown to be unstable during passage through the mouse GI tract. In a recent study, a maximum of 0.1% of the target population in the gut could be transduced^14^. Other studies have relied on bacteriophage λ, taking advantage of its temperate lifestyle. When infecting an *E. coli* cell, λ will either enter its lytic cycle and produce more virions, or enter lysogeny and integrate into the chromosome, thereby enabling the maintenance of transgenes such as a whole type I CRISPR-Cas system^20^ or dCas9^19^. Despite its ability to reproduce in the gut environment, previous work showed how λ fails to lysogenize the whole target population unless additional selection pressures are used^20^. Here we explore the use of engineered λ particles that employ receptors on the *E. coli* surface that are consistently expressed in the gut environment, enabling high delivery efficiencies.

The introduction of genetic modifications *in situ* further requires an efficient editing strategy. Cleavage of the bacterial chromosome by Cas9 leads to the death of most cells^21^. When combined with an efficient delivery strategy, this can be used in the development of sequence specific antimicrobials^13,22,23^. In recent work employing M13 as a delivery vector, antibiotics were used to kill target bacteria that did not receive the DNA payload and select those that survived Cas9 cleavage^14^. These bacteria will typically carry large uncontrolled deletions at the target position. While this strategy enables to perform *in situ* genetic modifications, it does so with very poor control and at the cost of imposing strong perturbations on the ecosystem. To achieve efficient editing without killing the target bacteria we turned to base editors. Base editors convert one base pair to another at a target locus without introducing a double-strand DNA break^24^ and have successfully been used in a broad range of bacterial species^25–32^.

Another desired feature of an *in situ* targeted mutagenesis strategy is that it should not spread transgenes. To achieve this we developed a DNA payload that harnesses the replication machinery of a phage-inducible chromosomal island (PICI)^33^. Our design ensures that the delivered DNA will not be replicated in recipient bacteria, while still allowing sufficient expression of the base editor to achieve efficient editing. This strategy enables the introduction of stable genetic perturbations to the majority of an *E. coli* population colonizing the mouse gut, without the need for a selection pressure or the maintenance of a transgene.

## Results

### Engineering of an efficient and selective DNA delivery vector for *E. coli* colonizing the mouse gut

The adsorption of phage Ur-λ to *E. coli* cells is determined by two main components of the capsid. First, the side tail fiber (*stf*) gene encodes for long appendages anchored at the base plate promoting reversible adsorption of the phage to target bacteria through interaction with the OmpC outer membrane porin^34,35^ (**Fig. 1A**). Second, the tail tip protein gpJ recognizes the LamB outer membrane porin and results in an irreversible binding of the phage to the cell surface^36,37^.

**Figure 1:**
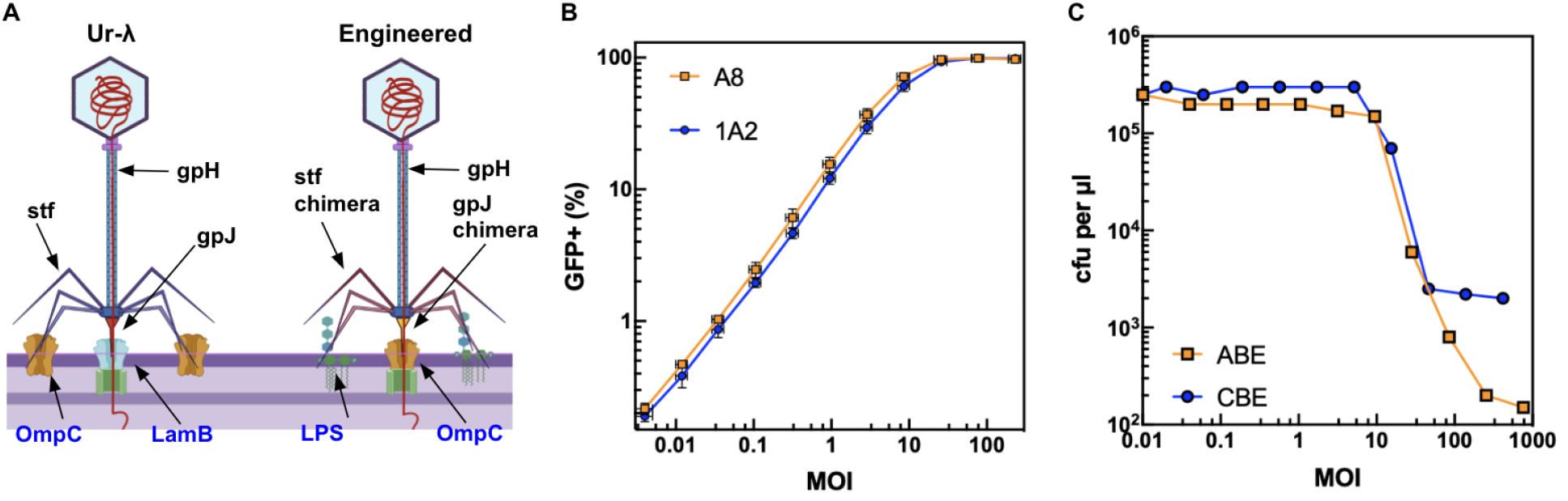
Engineering of an efficient and selective DNA delivery vector for *E. coli* colonizing the mouse gut. **A)** Schematic of the Ur-λ phage injection mechanism. Left panel: The adsorption of phage Ur-λ to *E. coli* cells is determined by three main components of the capsid. The side tail fiber (stf) gene encodes for long appendages anchored at the base plate, which promote adsorption of the phage to target bacteria through interaction with the OmpC outer membrane porin. The tail tip protein gpJ recognizes the LamB outer membrane porin. After binding of gpJ to its receptor, the gpH protein forms a tube through the periplasm and DNA is injected into the cytoplasm of the bacteria. Right panel: Engineered λ-derived particles with λ-P2 STF chimeras recognizing the LPS and with gpJ chimeras recognizing OmpC. **B)** Delivery efficiency of gpJ cosmid variants (1A2, orange line; A8, blue line) with a payload encoding *sfGFP* (plasmid p513) into *E. coli* s14269, measured by flow cytometry (excitation: 488 nm, emission: 530/30 BP). X axis: Multiplicity of injection (MOI; ratio of packaged cosmids to bacteria). Y axis: Percentage of GFP+ population after a 45-minute incubation. The graph shows the average and standard deviation of an experiment performed in triplicate. **C)** MOI-dependent adenine (ABE, p1396) and cytosine base editing (CBE, p2327) of β-lactamase on the strain *MG1655-bla*. Y axis: Colony-forming units (cfu) per μl on carbenicillin plates.

The adaptation of *E. coli* K-12 to the mouse gut selects for mutants that downregulate the expression of the maltose operon, which includes *lamB*, and such mutants being resistant to bacteriophage λ^38,39^. To circumvent this issue and ensure high and consistent delivery efficiencies, we engineered a variant of the Ur-λ capsid that uses a different receptor, the outer membrane porin OmpC. The expression of OmpC is upregulated in high-osmotic conditions typical of the gut and is known to be important for growth in the gut environment^40,41^. OmpC is also commonly used as a primary receptor by phages, including T4 and some lambdoids^42,43^.

We constructed different gpJ chimeras by fusing the C-terminal portion of naturally occurring gpJ variants found in *E. coli* phages to the λ *gpJ* gene. Additionally, to avoid competition between the new gpJ variants and the natural Ur-λ STF for binding to the same receptor, OmpC, we further constructed a chimera between the N-terminal part of Ur-λ STF and the C-terminal part of the phage P2 tail fiber, known to recognize the lipopolysaccharides of *E. coli* K-12 strains^44^. Chimeric gpJ variants were integrated in the Ur-λ prophage genome while the *stf* gene was removed and complemented by the P2-STF chimera encoded on a plasmid (p938). The Ur-λ prophage further carries the CI857 mutation making it heat inducible, and has its *cos* site inactivated, enabling the packaging of cosmids present in the cell while no phage DNA is being packaged^45^.

To evaluate capsid variants we packaged cosmids designed to express a *sfGFP* gene (plasmid p513), and measured delivery efficiency in different strains by flow cytometry after transduction^45^. A strain deleted for the *lamB* gene was used as recipient to identify gpJ variants unable to bind to LamB. In addition, to easily screen gpJ chimeras against two natural OmpC variants (from *E. coli* MG1655 or *E. coli* EDL933), we deleted *ompC* and expressed it from a plasmid (p1471 and p1472, respectively). One of the gpJ chimeras, A8, recognizes the OmpC receptor present in both the wild-type MG1655 and EDL933 strains; another one, 1A2, recognizes the OmpC receptor of *E. coli* EDL933 only (**Fig. S1A-C**). The functionality of the chimeric λ-P2 STF was further confirmed by showing markedly improved delivery efficiencies compared to capsids lacking an STF (**Fig. S1D**). Both gpJ variants enabled to transduce more than 90% of a target bacterial population at a multiplicity of injection (MOI) of ~20 (**Fig. 1B**).

Subsequently, we constructed cosmids carrying either an adenine base editor (ABE = ABE8e)^46^ or a cytosine base editor (CBE = evoAPOBEC1-nCas9-UGI)^47^. We first established that after the transformation of MG1655, these cosmids could efficiently edit two different targets. The first target was the active site triad of mCherry (M71, Y72, G73)^48^. Plasmid p2325 (ABE) was programmed to introduce the mutations M71T and Y72H while plasmid p2326 (CBE) was programmed to introduce the mutations G73N or G73D. For the second target, the β-lactamase (*bla*) gene was inserted in the *wbbL* locus of MG1655 (MG1655-*bla*); plasmid p1396 (ABE) was programmed to introduce the active site mutation K71E or K71R^49,50^ and plasmid p2327 (CBE) was programmed to introduce a premature stop codon (Q37*), resulting in the strain’s re-sensitization to carbenicillin. Both adenine and cytosine edits were obtained for both targets with >99% efficiency (**Fig. S2**).

We then investigated the feasibility of delivering the base editors encoded in plasmids p1396 (ABE) and p2327 (CBE) with our engineered λ particles to efficiently edit the target *bla* gene in MG1655-*bla* without selecting for the transduction of the cosmid (**Fig. 1C**). The λ-derived particles were produced and incubated with MG1655-*bla* for 2 hours at different MOIs. Base editing efficiency was measured by colony counting after overnight incubation on carbenicillin plates. ABE or CBE resulted in a ~10^4^ fold and ~10^3^ fold reduction of cell growth on carbenicillin plates at high MOI, respectively, showing that up to 99.99% of the bacterial population was edited and the β-lactamase gene inactivated. The reduction in plating efficiency at increasing MOIs is consistent with transduction rates observed in this experiment, with plating efficiency starting to drop as the majority of cells receive the cosmid (**Fig. S3**). Sanger sequencing of six colonies showed a base edit at position 7A in the editing window for all ABE samples, as well as bystander mutations at position 1A and 8A. Six sequenced CBE clones showed a single base pair change at position 4C in the editing window. Collectively, these experiments demonstrate the successful re-sensitization of a bacterial population to antibiotics by the use of base editors *in vitro*.

### Engineering of a non-replicative DNA cosmid

When considering the delivery of a DNA payload to edit a bacterial population in patients, it is highly desirable to avoid the dissemination of transgenes. To this end, we developed a cosmid that only replicates in the production strain and not in recipient bacteria, therefore preventing transfer of the DNA payload to any progeny cells. We modified our cosmid by replacing the p15A origin of replication with that of a phage-inducible chromosomal island (PICI) which requires a specific primase gene for replication^51^. We constructed a production strain expressing the primase gene on an additional plasmid under the control of the 2,4-diacetylphloroglucinol (DAPG)-inducible PhlF promoter^52^ (plasmid p1321) (**Fig. 2A**).

**Figure 2:**
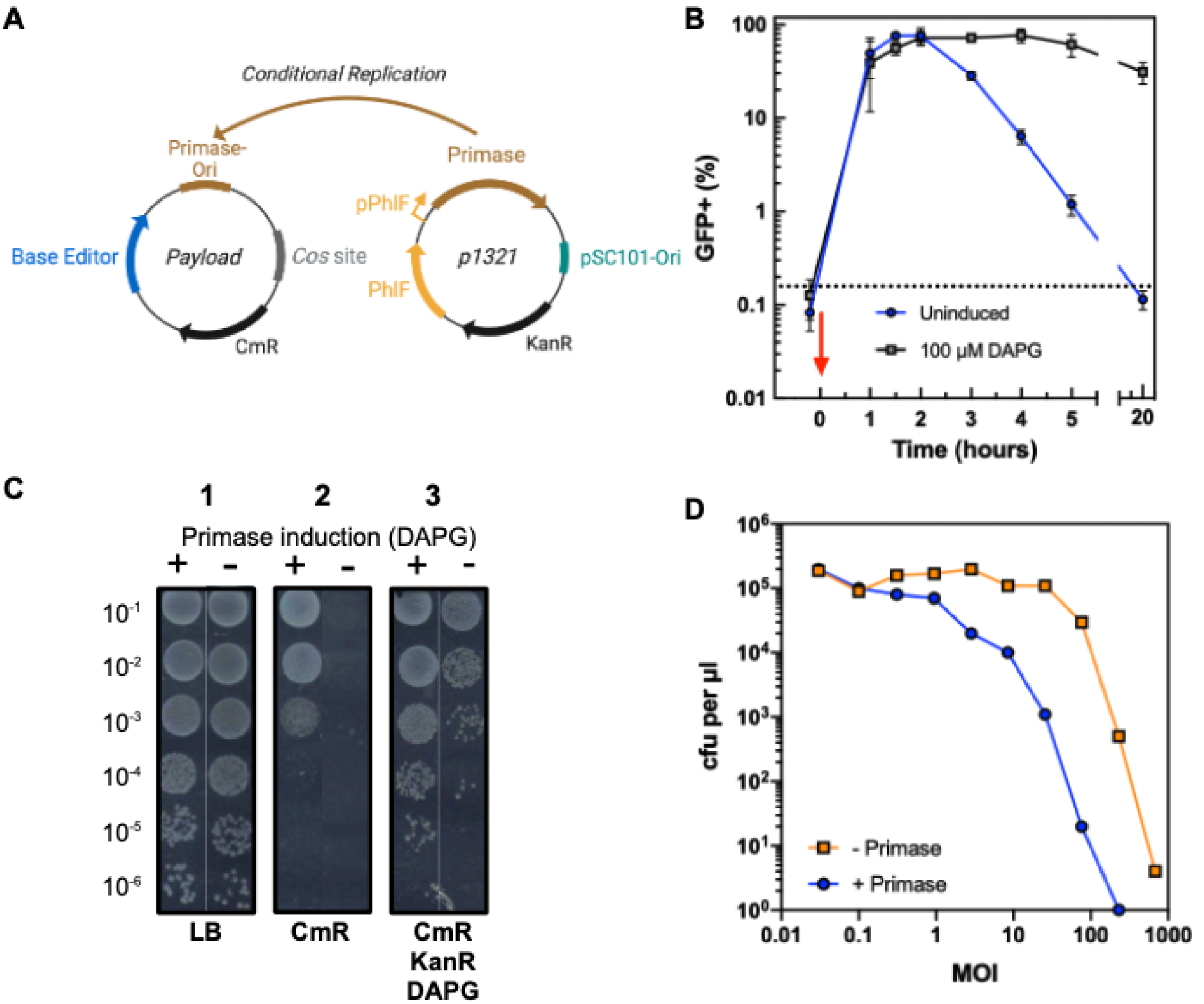
A non-replicative DNA payload can efficiently edit a target bacterial population. **A)** Schematics for the conditional replication of a cosmid in an *E. coli* production strain. Plasmid replication requires both the primase protein and the primase origin of replication. Upon delivery to recipient cells the cosmid cannot replicate in the absence of the primase. **B)** Plasmid stability was investigated *in vitro* with a time-course assay. Bacteria carrying an inducible primase plasmid without inducer (blue line), or a primase plasmid with 100 μM DAPG inducer (black line) were delivered with a cosmid harboring a *sfGFP* gene at MOI ~40. Dashed line: Background fluorescence of cells before transduction. The red arrow depicts the time at which the λ cosmid was added to the cells. Samples were taken at different time points and analyzed in a flow cytometer (excitation: 488 nm, emission: 530/30 BP; Attune NxT Thermo Scientific). The graph shows the average and standard deviation of an experiment performed in triplicate. **C)** Serially diluted cells carrying an induced (+) or uninduced (-) primase plasmid were plated 5 hours after transduction with a payload carrying the conditional origin of replication on LB-agar (1), LB-agar supplemented with chloramphenicol; CmR 25 μg ml^-1^ (2), or LB-agar supplemented with kanamycin; KanR 50 μg ml^-1^, chloramphenicol 25 μg ml^-1^, and DAPG 100 μM (3). **D)** Adenine base editing of β-lactamase on the *E. coli* genome after cosmid transduction *in vitro* using the non-replicative primase payload. A MG1655 strain encoding the β-lactamase gene was transduced in the presence (blue line) or absence (orange line) of the primase protein expressed inside the cell. Transduced cells were plated on LB with or without carbenicillin 2 hours post transduction at different MOIs and base editing efficiency was analyzed via colony counting the following day.

A time-course experiment was performed to investigate the plasmid stability in the presence or absence of the primase protein (**Fig. 2B**). The stability was measured by flow cytometry over time after transduction in *E. coli* MG1655 + p1321 using packaged Ur-λ cosmids encoding a *sfGFP* reporter gene in presence or absence of the inducer DAPG. After transduction, the fluorescence signal increased over time and reached a maximum within 1-2 hours in both conditions; after this time point, the GFP signal started decreasing for the bacteria without primase, while it remained constant for the induced cells. After 5 hours of incubation, only ~1% of the cells were positive for GFP in the absence of primase, compared to ~75% of the induced cells. These results demonstrate that the payload does not replicate in the absence of a primase protein. The absence of replication was further confirmed by plating on LB agar supplemented with chloramphenicol five hours after transduction. No colonies appeared with the uninduced-primase sample in the absence of DAPG. In contrast, the primase-induced sample grew on chloramphenicol-supplemented plates (**Fig. 2C**).

Our next focus was on determining whether the transient expression of a base editor by a non-replicative DNA payload would be sufficient to edit a whole target bacterial population. We constructed a conditionally replicative cosmid carrying the ABE programmed to target the *bla* gene (p2328), and packaged it in the engineered λ capsid with 1A2 gpJ and λ-P2 STF chimera. The packaged cosmid was used to transduce *E. coli* MG1655 carrying the EDL933 OmpC receptor and the *bla* gene in the presence or absence of the expressed primase protein (*s14269-bla*). Cells were then plated with or without carbenicillin and base editing efficiency was analyzed via colony counting the following day (**Fig. 2D**). While editing efficiency was slightly reduced in the absence of plasmid replication, the ABE still resulted in a ~10^4^ fold reduction of cell growth on carbenicillin at MOIs >100 using the non-replicative payload. This result demonstrates that up to 99.99% of the bacterial population was edited and that the β-lactamase gene was successfully inactivated.

### Targeted base editing of *E. coli* in the mouse gut using a non-replicative cosmid

We generated a streptomycin-resistant variant (*rpsLK42R*) of the s14269-*bla* strain (s21052) that could be used in an *in vivo* mouse colonization model. BALB/c mice were treated with streptomycin to allow for the intestinal engraftment of orally administered *E. coli* s21052. Five days after colonization, mice were orally gavaged with purified packaged cosmids equipped with gpJ 1A2 and the λ-P2 STF chimera, and an ABE targeting the *bla* gene (p2328) **(Fig. 3A**). To quantitatively monitor the base editing efficiency in mouse stool samples, we established a digital droplet PCR (ddPCR) assay based on two competing probes with Locked Nucleic Acid bases and labeled with either HEX or FAM dyes. The two probes were designed to target either the wild-type or the base-edited sequence, enabling the relative quantification of both genotypes from mouse stools (**Fig. S4** and methods).

**Figure 3:**
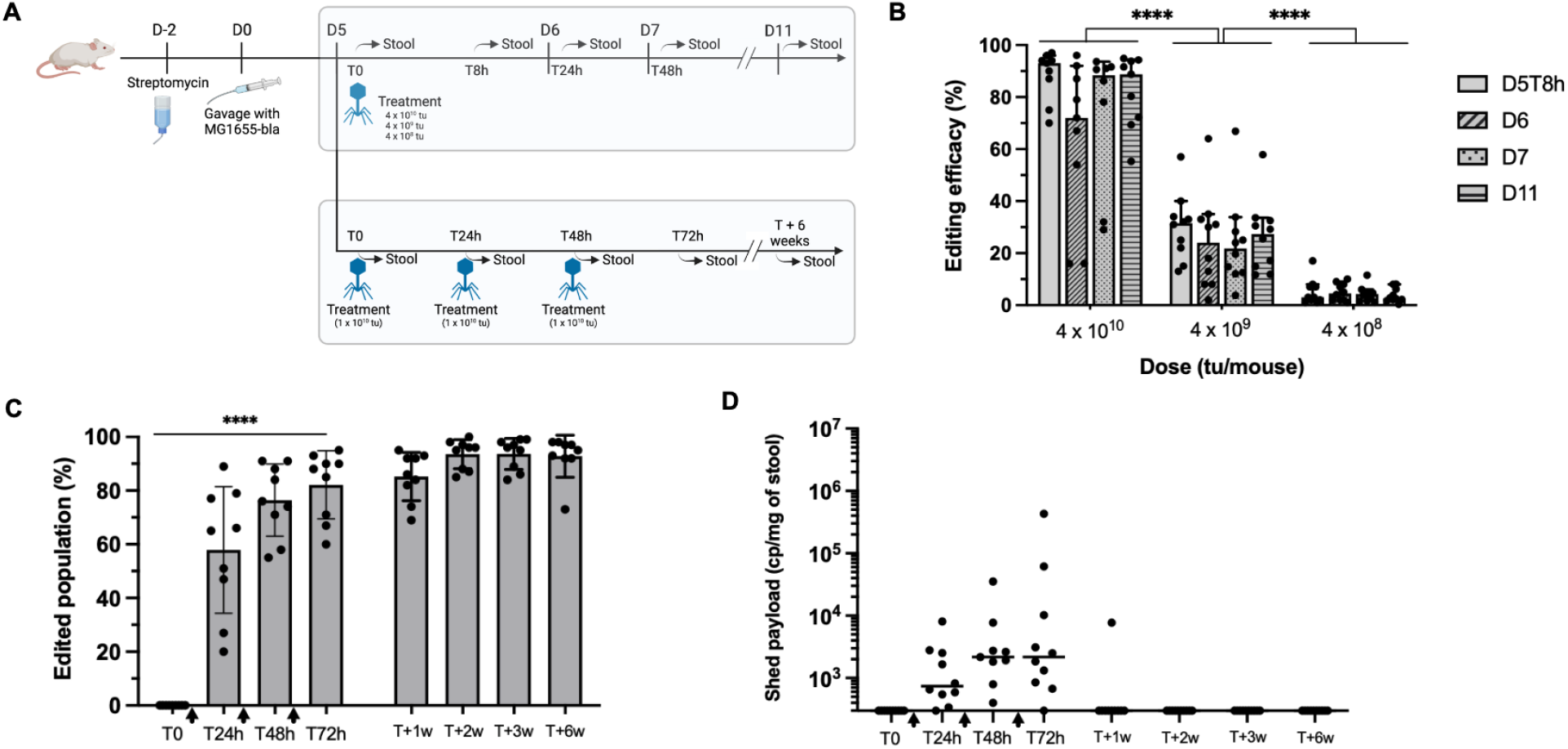
Targeted adenine base editing on the *E. coli* genome in gut of BALB/c mice after packaged λ cosmid treatment using a non-replicative payload. **A)** Summary of the experimental setup. In one arm we investigate the dose-response and in the other the impact of multiple doses on the treatment efficacy. **B)** Editing efficiency at different time points for a single dose with increasing concentrations. Points show individual mice, bars indicate the median, and error bars the standard deviation (**** p<0.0001 by ANOVA with Tukey’s multiple comparison test). **C)** Editing efficiency after multiple treatments (**** p<0.0001 by RM-ANOVA, with t-test for linear trend; treatments are indicated with black arrows on the x-axis). **D)** Number of copies of payload recovered in the stool and quantified by ddPCR. Bars represent the group median, with 95% confidence interval where applicable.

Larger doses of packaged cosmids yielded increased base editing efficacy of the total *E. coli* population present in the mouse gut, with a maximum median efficacy of 93% at the highest dose (4 x 10^10^ particles), as soon as 8 hours after treatment (**Fig. 3B**). This editing efficacy was calculated from direct quantification of target DNA molecules in stool samples frozen at the indicated collection time, rather than by repatching on agar plates, which would require overnight incubation. This ensured that the reported numbers were a measure of the editing efficacy at sampling time. The median editing efficacy was reduced to 32% and 3% with doses of 4 x 10^9^ and 4 x 10^8^ particles, respectively. Additionally, repatching colonies onto a selective medium was performed to corroborate ddPCR data (**Fig. S5**). Importantly, we observed that edited populations remained stable for at least six days after treatment, suggesting no obvious fitness cost of the targeted genetic modification. This demonstrates that our base editing approach is capable of inducing stable modification of bacterial genes in the gut microbiome.

We then assessed whether administering several treatments would increase the relative abundance of the edited population. We selected an intermediate dose (1 x 10^10^ particles), and administered 1 dose per day, for 3 consecutive days to the mice. Each dose successfully increased the median editing efficacy from 65% to 76%, and finally 88% of the target bacterial population. While the total *E. coli* s21052 colonization levels decreased over the 6-week experiment, the average proportion of edited bacteria remained stable until the end, showing the absence of fitness cost as well as the stability of the genetic modification (**Fig. 3C and S6**).

Over-time, the proportion of edited versus non-edited bacteria was seen to slightly fluctuate in feces pellets recovered from individual mice. The highest ratio was measured at the 3-week time point in a mouse, where 99.7% of the bacterial population carried the desired modification as measured by ddPCR. We did not detect any excreted payload in the stool of treated animals 11 days after the last treatment, and only detected the payload in 1 out of 10 animals 4 days after the last treatment (**Fig. 3D**). This showed that the non-replicative payload was not maintained in the target bacteria, while still allowing for efficient editing of the bacterial population.

## Discussion

In this study, we demonstrate the efficient and durable genetic modification of a bacterial population in the gut environment after the administration of non-replicative phage-based delivery particles equipped with base editors. While previous studies have demonstrated DNA delivery of a transgene to bacteria colonizing the mouse gut, it remains a challenge to ensure that the majority of the target population receives a DNA payload. Selection pressures such as the administration of antibiotics^14^ or lytic phages have thus been used to kill bacteria that did not receive the payload^20^. Here we show that by carefully designing a phage vector to effectively target *E. coli* in the gut environment, it is possible to deliver DNA to the vast majority of the bacterial population without resorting to selection.

Our engineering efforts focused on the construction of λ particles with chimeric side tail fibers (STF) and tail tip (gpJ) proteins. The specificity of STF was changed from OmpC to the LPS, while that of GpJ was changed from LamB to OmpC. These modifications ensure that our vector can recognize surface determinants consistently expressed by a specific *E. coli* target strain in the gut environment. We anticipate that the modular swapping of receptor binding domains demonstrated here will allow targeting various strains of *E. coli* as well as other species with engineered λ particles.

Future therapeutic applications of *in situ* base editing will be greatly facilitated if the dissemination of the payload is limited as much as possible. For this reason, we explored the possibility of delivering DNA payloads that do not replicate in recipient bacteria. To obtain a conditionally replicating cosmid, we addressed several requirements during our design process. The replication of our cosmid should be conditional in the presence of a protein that can be expressed in the donor bacteria, but is absent or rare in recipient cells. This origin of replication should further enable the efficient replication of the cosmid DNA in concatemeric form, which is required for efficient packaging into the capsid of bacteriophage λ^54,55^. Natural parasites of *Siphovirus* bacteriophages, known as PICIs, package themselves into a helper phage capsid^51^ and were used in the design. The origin of replication from PICIs requires a primase protein, whose gene is found in 1,632 out of 25,759 (6.3%) *E. coli* genome assemblies from RefSeq (as of Nov 25^th^ 2021). This sequence is also rare in other Enterobacteriales (708 of 64,071 genomes - 1.1%). The primase gene is not constitutively expressed in bacteria as it is present as part of a PICI; hence, its expression is dependent on induction of the PICI. Therefore, replication of the payload is unlikely, even in strains that contain a primase gene. This approach also ensures that the edited strain has no reduced fitness due to payload maintenance, which could lead to the loss of the edited strain in complex communities of the gut microbiome. While efficient base editing was previously demonstrated in bacteria^56,57^, this was performed under the prolonged expression of the base editor from a replicative plasmid maintained in the bacteria with antibiotic pressure. Here we could show that the transient expression of a base editor from a non-replicative plasmid without the need for antibiotic selection is sufficient to edit target bacteria.

We have shown that the inactivation of a β-lactamase gene incurred no fitness cost in the gut environment of mice, as the proportion of edited cells remained stable for at least 6 weeks. Measuring how the relative abundance of bacteria edited with this approach changes over time is an exciting avenue to investigate the genetic determinants of bacterial fitness in the native gut environment.

For therapeutic applications, the effect of the edits on the bacteria’s fitness will be a critical consideration. In the absence of selection pressure to eliminate non-edited bacteria, or when a delivery vector does not achieve full penetrance, a small fraction of unedited bacteria will remain. If the edit is costly, unedited strains will take over the population. Restoration of the wild-type bacteria would for multiple diseases result in return of the disease phenotype. However, one disease area where this problem might be mitigated is when bacterial antigens mimic human antigens and drive disease. Recent research has highlighted the importance of bacterial peptide mimics in autoimmune and inflammatory diseases^6,7^, cancers^8^, as well as immunotherapies^1^. The properties of the peptide mimic are typically independent of the function carried out by the bacterial protein. It should therefore be possible to eliminate or create peptide mimics, while preserving functionality of the bacterial protein. Absence of fitness cost may facilitate the durability of the responses, which is of particular importance in chronic disease.

While some modification of interest might be out of reach of base editors, we envision that the platform can be easily adapted to other gene editors such as prime editors^58^, or RNA-guided transposons^59–62^, and more. Payload size limitations as reported for mammalian delivery systems such as adeno-associated viruses are not a concern here as λ virions naturally package 48.5kb^63^. Future work will focus on the engineering of both the delivery vectors for bacterial species of interest, and the appropriate gene editing strategy. Our strategy provides novel opportunities to investigate bacteria of the microbiome, while opening up a wide array of therapeutic options for altering disease-driving genes of resident or pathogenic bacteria not easily targeted by conventional methods.

## Methods

### Strains and media

The *E. coli* and λ prophage genomes were engineered using a strategy relying on λ red homologous recombination coupled to Cas9 targeting^64,65^. Packaged cosmids were produced using an engineered *E. coli* K-12 strain carrying the thermosensitive cI857 λ prophage with its *cos* site deleted. When needed, a constitutively expressed SrpR repressor^52^ was inserted in the *lacZ* locus, in order to repress the expression of genes in the production strain. The chimeric λ-P2 *stf* gene was expressed in a plasmid *in trans* (p938), with the *stf* gene deleted from the prophage. All experiments were performed with cells grown in LB plus 5 mM CaCl_2_, supplemented with antibiotics when necessary unless stated otherwise (chloramphenicol (12.5 to 25 μg ml^-1^), kanamycin (25 to 50 μg ml^-1^), trimethoprim (5 to 10 μg ml^-1^), streptomycin (50 to 100 μg ml^-1^), ampicillin (50 to 100 μg ml^-1^), and carbenicillin (50 to 100 μg ml^-1^)). 2,4-Diacetylphloroglucinol (DAPG; 100 μM, Santa Cruz Biotechnology) was added to the media or plates in order to induce the pPhlF promoter^66^.

All titration assays were carried out with an *E. coli* MG1655 strain (NCBI no. NC_000913) modified to encode the EDL933 OmpC variant, if necessary (s14269), and carrying a plasmid expressing the primase gene (p1321) if needed. mCherry (GenBank no. QQM12952) and β-lactamase (GenBank no. ANG10794) were used as reporter proteins encoded on the *E. coli* MG1655 genome (MG1655-*mCherry* and MG1655-*bla*). For the chimeric gpJ/receptor analysis, the plasmid stability assays, and the delivery efficiency assays, superfolder Green Fluorescent Protein (sfGFP; GenBank no. AYN72676) was used as a reporter. *E. coli* MG1655 strains carrying the OmpC EDL933 receptor and the *bla* gene were used for the primase experiments (s14269 and MG1655-*bla*). Streptomycin-resistant mutants of *E. coli* s14269 were used for *in vivo* experiments (s21052). Strain genotypes are listed in **Table S1.**

### Cloning and plasmid construction

Standard DNA cloning was performed with chemically competent DH10B cells (Thermo Scientific) or a modified K-12 strain carrying the SrpR repressor^52^. Molecular cloning was carried out using Gibson Assembly^67^. Base editors ABE8e^46^ and evoAPOBEC1-nCas9-UGI^47^ were codon-optimized for *E. coli* and synthesized (Twist Bioscience). The obtained DNA fragments were assembled and cloned downstream of the pSrpR promoter for all constructs. Base editor expression levels were tuned using DNA libraries of ribosomal binding sites^68^. Sequences for the λ-P2 STF chimera were amplified from phage P2 and phage λ genomic DNA and cloned downstream of the inducible pPhlF promoter. The primase gene from the *E. coli* CFT073 strain (NCBI no. AAN79964, locus AE016759_238) was amplified from the CFT073 genome and cloned into plasmids p2076 or p1321 (constitutive or DAPG-inducible promoters, respectively). The cohesive end site (*cos*) of the λ genome was cloned onto the p15A payload to enable packaging into λ cosmid particles. All plasmids were purified using a Plasmid DNA Miniprep Kit (Omega Bio-Tek) and sequence-verified by Sanger sequencing (Eurofins Genomics). Guide RNAs and oligonucleotide sequences are listed in **Tables S2-S4**. A plasmid map of the non-replicative cosmid encoding the adenine base editor and a guide RNA is depicted in **Fig. S7**.

### Packaged cosmid production

Packaged cosmid production was performed with an engineered *E. coli* K-12 strain derived from CY2120^45^ carrying a modified λ prophage (CY-1A2, CY-A8, or CY-Ur-λ, **Table S1**). For the packaged cosmids in Fig. 1B, the production strains contained the plasmids p513 and p938; and for the packaged cosmids in Fig. 1C and Fig. S3, the production strains carried the plasmids p1396 and p938, or p2327 and p938. For the packaged cosmids in Fig. 2B and 2C, the production strains contained the plasmids p1324 and p1321; for the packaged cosmids in Fig. 2D and Fig. 3, the production strains carried the plasmids p2328, p938 and p2076; and for the packaged cosmids in Fig. S1, the production strains contained the plasmid p513, as well as the plasmid p938 (λ-P2 STF chimera) when indicated.

Cultures were performed in LB medium supplemented with 5 mM CaCl_2_ with the appropriate antibiotics and 100 μM DAPG if necessary in order to induce λ-P2 STF expression or primase expression. Production strains were grown overnight at 30°C in liquid media in an orbital shaker, diluted 1:6 the next day in fresh media supplemented with antibiotics and DAPG when needed, and grown for 30 minutes at 30°C. Packaged cosmid production was heat-induced at 42°C for 45 minutes. After that, cell cultures were shifted to 37°C for 3 to 6 hours in an orbital shaker at 180 rpm (New Brunswick Innova 44). Samples were centrifuged for 10 min at 4,500g and cell pellets were resuspended and lysed with B-PER reagent (Thermo Scientific) (1/10 of the initial volume of cosmid production for *in vitro* assays and 1/50 of the initial volume of production for *in vivo* assays) and lysozyme (100 μg ml^-1^; Applichem Lifescience). DENARASE (10,000x; c-Lecta) was added to the reaction to degrade residual DNA and RNA at a dilution of 1:10,000. Bio-Beads (SM-2 resin; Bio-Rad) were added to the lysis reaction and samples were incubated for 1 hour in a mini-shaker (PS-3D, Grant-Bio) at room temperature. Samples were centrifuged for 5 min at 16,000g, and supernatants were sterile filtered (0.22 μm pore size; Sartorius Minisart). For *in vivo* administration, cosmids were concentrated and buffer-exchanged against Phosphate Buffered Saline (PBS) by tangential flow filtration (MWCO 100 kDA; Sartorius Vivaflow 200). Packaged cosmid concentration was analyzed by *E. coli* transduction with diluted cosmid stocks (1:10 dilutions) and consecutive colony counting on chloramphenicol plates after overnight incubation at 37°C.

### Packaged cosmid delivery efficiency

Delivery efficiency was analyzed using cosmid particles equipped with λ-P2 STF chimera and gpJ 1A2 or A8 into the strain s14269. The cosmid encodes a *sfGFP* gene under a constitutive promoter. The cells were grown in LB supplemented with 5 mM of CaCl_2_ to an OD_600_ of 0.2 to 0.6. Cell density was adjusted to OD_600_ = 0.025 in fresh LB supplemented with 5 mM CaCl_2_, and 90 μl of cell culture was mixed with 10 μl of each cosmid serially diluted in LB plus 5 mM CaCl_2_ (1:3 dilutions) to reach different MOIs. The samples were incubated for 45 min at 37°C and 8 μl was added to 250 μl ice-cold PBS plus 1 mg ml^-1^ kanamycin prior to analysis by flow cytometry (excitation: 488 nm, emission: 530/30 BP; Attune NxT Thermo Scientific).

### Plasmid stability assay

Plasmid stability was investigated *in vitro* with a time-course assay. *E. coli* MG1655 carrying a DAPG-inducible primase plasmid (p1321) with or without 100 μM DAPG were grown to an OD_600_ of 0.2 to 0.6 in LB plus 5 mM CaCl_2_ and 50 μg ml^-1^ kanamycin. Samples were then diluted to an OD_600_ of 0.01 in fresh LB plus 5 mM CaCl_2_ and 50 μg ml^-1^ kanamycin plus/minus 100 μM DAPG, treated with a packaged cosmid harboring the *sfGFP* gene and the conditional primase origin of replication (p1324) at an MOI ~40, and subsequently incubated in an orbital shaker at 37°C. 1-5 μl samples were taken at different time points, mixed with 250 μl ice-cold PBS supplemented with 1 mg ml^-1^ kanamycin, and analyzed in a flow cytometer (excitation: 488 nm, emission: 530/30 BP; Attune NxT Thermo Scientific). To maintain the cells in the exponential growth phase, the samples were diluted 1:5 into fresh LB media supplemented with 50 μg ml^-1^ kanamycin plus/minus 100 μM DAPG every 2 hours.

### Base editing *in vitro*

The *E. coli* MG1655-*mCherry* strain was transformed with the base editor payloads p2316 or p2326, grown for 2 hours in SOC medium (30°C, 180 rpm), and selected on chloramphenicol plates overnight at 30°C. Forty-eight individual colonies were resuspended in 250 μl Phosphate Buffered Saline supplemented with 1 mg ml^-1^ kanamycin in a 96-well plate and mCherry fluorescence was measured by flow cytometry (excitation: 561 nm, emission: 620/15 BP; Attune NxT Thermo Scientific). As a control, base editors were transformed in the absence of a guide RNA and the mCherry fluorescence of 2 colonies was analyzed.

The *E. coli* strain MG1655-*bla* transformed with the base editor payloads (p1396 and p2327) was grown for 2 hours in SOC medium (30°C, 180 rpm) prior to spotting of 10 μl of individual cell dilutions on chloramphenicol/carbenicillin plates, as well as on chloramphenicol plates. As a control, base editors were transformed in the absence of a guide RNA on the payload. Editing efficiency was analyzed by colony counting on plates after overnight incubation at 30°C.

Base editing of transduced packaged cosmids in *E. coli* strain MG1655-*bla* was performed similarly to transformation assays. The target strain was cultured to mid-log phase, diluted to an OD_600_ of 0.025 (Fig. 1C) or 0.005 (Fig. 2D), and transduced with serial dilutions (1:3) of the produced packaged cosmid in a 96-well plate. Cells were grown for 2 hours in LB medium supplemented with 5 mM CaCl_2_ (30°C, 180 rpm) prior to spotting of individual dilutions on carbenicillin plates. As a control, cells were treated with LB media instead of packaged cosmid solution. For all experiments, the target gene was amplified via PCR from the genome of individual colonies and base editing was confirmed by Sanger sequencing.

### Mouse experiments with *E. coli* colonization and packaged cosmid treatment

Specific pathogen-free 5 to 9 week old female BALB/cYJ mice were supplied by Charles River Laboratories and housed in an animal facility in accordance with Institut Pasteur guidelines and European recommendations. Animal procedures were approved by the Institut Pasteur (approval ID: 20040) and the French Research Ministry (APAFIS ID: 28717) and animal experiments were performed in compliance with applicable ethical regulations. Water and food were provided *ad libitum*, unless stated otherwise.

Animals were acclimated for 5 days before streptomycin sulfate (5 mg ml^-1^; Sigma-Aldrich S9137) was added to the autoclaved drinking water to decrease the number of facultative aerobic/anaerobic resident bacteria^69^. Drinking water containing streptomycin was prepared fresh weekly. Three days later (D0), mice were orally gavaged with approximately 1 x 10^8^ cfu of strain s21052, grown overnight in LB and resuspended in 200 μl of sterile gavage buffer (20% sucrose, 2.6% sodium bicarbonate, pH 8). Starting at D5, mice were orally administered (200 μl per mouse) with either gavage buffer or with packaged cosmids diluted 1:1 in buffer. The appropriate dose was achieved by diluting the packaged cosmid suspension in PBS, prior to formulation in buffer, and checked by *E. coli* transduction.

### Evaluation of edited *E. coli* from mouse feces by direct plating

Fresh fecal samples were collected at D0 and subsequent relevant time points as a proxy to assess intestinal colonization levels of s21052. Briefly, fecal samples were weighed on an analytical balance and 1 ml of PBS was added. Samples were incubated for 2 min at room temperature and suspended by manual mixing and vortexing. Serial dilutions were performed in PBS, and 5 μl of each dilution was spotted onto Drigalski agar plates (Bio-Rad) supplemented with 100 μg ml^-1^ of streptomycin, and plates were incubated overnight at 37°C. Estimation of editing efficacy was performed the following day by repatching individual colonies (up to 12 colonies per mouse and per time point) onto agar plates, with or without 50 μg ml^-1^ carbenicillin to investigate the loss of resistance to β-lactams subsequent to editing of the *bla* gene. Additionally, separate fecal samples were collected and frozen at −80°C within one hour of collection to assess the editing efficacy by ddPCR.

### Base editing quantification by droplet digital PCR

Primers F3 (5’-GGATCTCAACAGCGGTAAG-3’) and R3 (5’-GGCATCAACACGGGATAATA-3’), both with a melting temperature of 61°C, were designed to amplify a 112-bp region of the *bla* gene in *E. coli* s21052 spanning the target site for base editing. Two Taqman probes were designed to bind this amplicon with the target site towards the middle of the probes, before or after successful editing (**A** to **G**): P1 (5’-FAM-CT+TT+T**+A**+AA+GTT+C+T+GC-3’) and P2 (5’-HEX-CT+TT+T**+G**+AAGTT+CT+GC-3’). Each probe contained a different fluorophore (FAM or HEX), as well as carefully positioned Locked Nucleic Acid bases (LNA; symbolized by the base A, T, C, or G preceded by a “+” sign in the sequences above). LNA nucleotides allow for a greater Tm difference between matching and mismatching probes while retaining a small probe size, further improving discrimination^70^. Tm for either probe matching their specific sequence was predicted to be 66°C, compared to 55°C in case of binding to the non-matching sequence (OligoAnalyzer Tool, IDT).

Reactions were conducted in a 8-μl final volume, with PerfeCTa Multiplex qPCR Toughmix, 100 nM fluorescein, 250 nM of each primer and 250 nM of each probe, using a Naica ddPCR system (Stilla Technologies). The following 2-step cycling program was applied: initial denaturation for 3 minutes at 95°C, followed by 50 cycles of 95°C for 10 sec and 57°C for 30 seconds. This Taqman assay was validated for specificity using purified genomic DNA from overnight bacterial cultures of either wild-type s21052 or *in vitro*-edited s21052 (**Fig. S4**), and fluorescence spillover compensation was carried out using the appropriate control reactions following the manufacturer’s recommendations.

Stool samples collected from mice were weighed, resuspended at 100 mg ml^-1^ in ultrapure water, homogenized and heat-treated at 98°C for 10 minutes. After brief vortexing and a one-minute cooldown at room temperature, supernatant was pipetted from the top of the suspension to avoid major debris, diluted at least 10 times in ultrapure water, and analyzed immediately by ddPCR without further processing.

## References

1. Mager, L. F. et al. Microbiome-derived inosine modulates response to checkpoint inhibitor immunotherapy. Science 369, 1481–1489 (2020).

2. Matson, V. et al. The commensal microbiome is associated with anti–PD-1 efficacy in metastatic melanoma patients. Science 359, 104–108 (2018).

3. Gopalakrishnan, V. et al. Gut microbiome modulates response to anti-PD-1 immunotherapy in melanoma patients. Science 359, 97–103 (2018).

4. Routy, B. et al. Gut microbiome influences efficacy of PD-1-based immunotherapy against epithelial tumors. Science 359, 91–97 (2018).

5. McCulloch, J. A. et al. Intestinal microbiota signatures of clinical response and immune-related adverse events in melanoma patients treated with anti-PD-1. Nat. Med. 28, 545–556 (2022).

6. Greiling, T. M. et al. Commensal orthologs of the human autoantigen Ro60 as triggers of autoimmunity in lupus. Sci. Transl. Med. 10, (2018).

7. Gil-Cruz, C. et al. Microbiota-derived peptide mimics drive lethal inflammatory cardiomyopathy. Science 366, 881–886 (2019).

8. Cullin, N., Antunes, C. A., Straussman, R., Stein-Thoeringer, C. K. & Elinav, E. Microbiome and cancer. Cancer Cell 39, 1317–1341 (2021).

9. Wong, S. H. & Yu, J. Gut microbiota in colorectal cancer: mechanisms of action and clinical applications. Nat. Rev. Gastroenterol. Hepatol. 16, 690–704 (2019).

10. Rekdal, V. M., Bess, E. N., Bisanz, J. E., Turnbaugh, P. J. & Balskus, E. P. Discovery and inhibition of an interspecies gut bacterial pathway for Levodopa metabolism. Science 364, (2019).

11. Zimmermann, M., Zimmermann-Kogadeeva, M., Wegmann, R. & Goodman, A. L. Mapping human microbiome drug metabolism by gut bacteria and their genes. Nature 570, 462–467 (2019).

12. Quigley, E. M. M. & Gajula, P. Recent advances in modulating the microbiome. F1000Research 9, F1000 Faculty Rev-46 (2020).

13. Citorik, R. J., Mimee, M. & Lu, T. K. Sequence-specific antimicrobials using efficiently delivered RNA-guided nucleases. Nat Biotechnol 32, 1141–5 (2014).

14. Lam, K. N. et al. Phage-delivered CRISPR-Cas9 for strain-specific depletion and genomic deletions in the gut microbiome. Cell Rep. 37, (2021).

15. High-efficiency delivery of CRISPR-Cas9 by engineered probiotics enables precise microbiome editing. Mol. Syst. Biol. 17, e10335 (2021).

16. Ronda, C., Chen, S. P., Cabral, V., Yaung, S. J. & Wang, H. H. Metagenomic engineering of the mammalian gut microbiome in situ. Nat. Methods 16, 167–170 (2019).

17. Neil, K., Allard, N., Grenier, F., Burrus, V. & Rodrigue, S. Highly efficient gene transfer in the mouse gut microbiota is enabled by the Incl2 conjugative plasmid TP114. Commun. Biol. 3, 523 (2020).

18. Allard, N., Neil, K., Grenier, F. & Rodrigue, S. The Type IV Pilus of Plasmid TP114 Displays Adhesins Conferring Conjugation Specificity and Is Important for DNA Transfer in the Mouse Gut Microbiota. Microbiol. Spectr. 10, e02303–21 (2022).

19. Hsu, B. B. et al. In situ reprogramming of gut bacteria by oral delivery. Nat. Commun. 11, 5030 (2020).

20. Yosef, I., Manor, M., Kiro, R. & Qimron, U. Temperate and lytic bacteriophages programmed to sensitize and kill antibiotic-resistant bacteria. Proc. Natl. Acad. Sci. 112, 7267–7272 (2015).

21. Cui, L. & Bikard, D. Consequences of Cas9 cleavage in the chromosome of Escherichia coli. Nucleic Acids Res. 44, 4243–4251 (2016).

22. Bikard, D. et al. Exploiting CRISPR-Cas nucleases to produce sequence-specific antimicrobials. Nat Biotechnol 32, 1146–50 (2014).

23. Neil, K. et al. High-efficiency delivery of CRISPR-Cas9 by engineered probiotics enables precise microbiome editing. Mol. Syst. Biol. 17, e10335 (2021).

24. Anzalone, A. V., Koblan, L. W. & Liu, D. R. Genome editing with CRISPR–Cas nucleases, base editors, transposases and prime editors. Nat. Biotechnol. 38, 824–844 (2020).

25. Rodrigues, S. D. et al. Efficient CRISPR-mediated base editing in Agrobacterium spp. Proc. Natl. Acad. Sci. 118, e2013338118 (2021).

26. Yu, S. et al. CRISPR-dCas9 Mediated Cytosine Deaminase Base Editing in Bacillus subtilis. ACS Synth. Biol. 9, 1781–1789 (2020).

27. Chen, W. et al. CRISPR/Cas9-based Genome Editing in Pseudomonas aeruginosa and Cytidine Deaminase-Mediated Base Editing in Pseudomonas Species. iScience 6, 222–231 (2018).

28. Zhang, Y. et al. Programmable adenine deamination in bacteria using a Cas9–adenine-deaminase fusion. Chem. Sci. 11, 1657–1664 (2020).

29. Zheng, K. et al. Highly efficient base editing in bacteria using a Cas9-cytidine deaminase fusion. Commun. Biol. 1, 1–6 (2018).

30. Volke, D. C., Martino, R. A., Kozaeva, E., Smania, A. M. & Nikel, P. I. Modular (de)construction of complex bacterial phenotypes by CRISPR/nCas9-assisted, multiplex cytidine base-editing. Nat. Commun. 13, 3026 (2022).

31. Pan, M. et al. Genomic and epigenetic landscapes drive CRISPR-based genome editing in Bifidobacterium. Proc. Natl. Acad. Sci. U. S. A. 119, e2205068119 (2022).

32. Banno, S., Nishida, K., Arazoe, T., Mitsunobu, H. & Kondo, A. Deaminase-mediated multiplex genome editing in Escherichia coli. Nat. Microbiol. 3, 423–429 (2018).

33. Penadés, J. R. & Christie, G. E. The Phage-Inducible Chromosomal Islands: A Family of Highly Evolved Molecular Parasites. Annu. Rev. Virol. 2, 181–201 (2015).

34. Montag, D., Schwarz, H. & Henning, U. A component of the side tail fiber of Escherichia coli bacteriophage lambda can functionally replace the receptor-recognizing part of a long tail fiber protein of the unrelated bacteriophage T4. J. Bacteriol. 171, 4378–4384 (1989).

35. Hendrix, R. W. & Duda, R. L. Bacteriophage lambda PaPa: not the mother of all lambda phages. Science 258, 1145–1148 (1992).

36. Randall-Hazelbauer, L. & Schwartz, M. Isolation of the Bacteriophage Lambda Receptor from Escherichia coli. J. Bacteriol. 116, 1436–1446 (1973).

37. Schwartz, M. Reversible interaction between coliphage lambda and its receptor protein. J. Mol. Biol. 99, 185–201 (1975).

38. Paepe, M. D. et al. Carriage of λ Latent Virus Is Costly for Its Bacterial Host due to Frequent Reactivation in Monoxenic Mouse Intestine. PLOS Genet. 12, e1005861 (2016).

39. Paepe, M. D. et al. Trade-Off between Bile Resistance and Nutritional Competence Drives Escherichia coli Diversification in the Mouse Gut. PLOS Genet. 7, e1002107 (2011).

40. Yoshida, T., Qin, L., Egger, L. A. & Inouye, M. Transcription Regulation of ompF and ompC by a Single Transcription Factor, OmpR*. J. Biol. Chem. 281, 17114–17123 (2006).

41. Warr, A. R. et al. Transposon-insertion sequencing screens unveil requirements for EHEC growth and intestinal colonization. PLOS Pathog. 15, e1007652 (2019).

42. Hantke, K. Major outer membrane proteins of E. coli K12 serve as receptors for the phages T2 (protein Ia) and 434 (protein Ib). Mol. Gen. Genet. MGG 164, 131–135 (1978).

43. Yu, F. & Mizushima, S. Roles of lipopolysaccharide and outer membrane protein OmpC of Escherichia coli K-12 in the receptor function for bacteriophage T4. J. Bacteriol. 151, 718–722 (1982).

44. North, O. I. & Davidson, A. R. Phage Proteins Required for Tail Fiber Assembly Also Bind Specifically to the Surface of Host Bacterial Strains. J. Bacteriol. 203, e00406–20 (2021).

45. Cronan, J. E. Improved plasmid-based system for fully regulated off-to-on gene expression in Escherichia coli: Application to production of toxic proteins. Plasmid 69, 81–89 (2013).

46. Richter, M. F. et al. Phage-assisted evolution of an adenine base editor with improved Cas domain compatibility and activity. Nat. Biotechnol. 38, 883–891 (2020).

47. Thuronyi, B. W. et al. Continuous evolution of base editors with expanded target compatibility and improved activity. Nat. Biotechnol. 37, 1070–1079 (2019).

48. Shaner, N. C. et al. Improved monomeric red, orange and yellow fluorescent proteins derived from Discosoma sp. red fluorescent protein. Nat. Biotechnol. 22, 1567–1572 (2004).

49. Tremblay, L. W., Xu, H. & Blanchard, J. S. Structures of the Michaelis complex (1.2 Å) and the covalent acyl intermediate (2.0 Å) of cefamandole bound in the active sites of the Mycobacterium tuberculosis β-lactamase K73A and E166A mutants. Biochemistry 49, 9685–9687 (2010).

50. Lietz, E. J., Truher, H., Kahn, D., Hokenson, M. J. & Fink, A. L. Lysine-73 is involved in the acylation and deacylation of beta-lactamase. Biochemistry 39, 4971–4981 (2000).

51. Fillol-Salom, A. et al. Phage-inducible chromosomal islands are ubiquitous within the bacterial universe. ISME J. 12, 2114 (2018).

52. Stanton, B. C. et al. Genomic Mining of Prokaryotic Repressors for Orthogonal Logic Gates. Nat. Chem. Biol. 10, 99–105 (2014).

53. Pédelacq, J.-D., Cabantous, S., Tran, T., Terwilliger, T. C. & Waldo, G. S. Engineering and characterization of a superfolder green fluorescent protein. Nat. Biotechnol. 24, 79–88 (2006).

54. Dawson, P., Hohn, B., Hohn, T. & Skalka, A. Functional empty capsid precursors produced by lambda mutant defective for late lambda DNA replication. J. Virol. 17, 576–583 (1976).

55. Enquist, L. W. & Skalka, A. Replication of bacteriophage λ DNA dependent on the function of host and viral genes: I. Interaction of red, gam and rec. J. Mol. Biol. 75, 185–212 (1973).

56. Gaudelli, N. M. et al. Programmable base editing of A·T to G·C in genomic DNA without DNA cleavage. Nature 551, 464–471 (2017).

57. Komor, A. C., Kim, Y. B., Packer, M. S., Zuris, J. A. & Liu, D. R. Programmable editing of a target base in genomic DNA without double-stranded DNA cleavage. Nature 533, 420–424 (2016).

58. Anzalone, A. V. et al. Search-and-replace genome editing without double-strand breaks or donor DNA. Nature 1–1 (2019) doi:10.1038/s41586-019-1711-4.

59. Klompe, S. E., Vo, P. L. H., Halpin-Healy, T. S. & Sternberg, S. H. Transposon-encoded CRISPR-Cas systems direct RNA-guided DNA integration. Nature 571, 219–225 (2019).

60. Strecker, J. et al. RNA-guided DNA insertion with CRISPR-associated transposases. Science 365, 48–53 (2019).

61. Vo, P. L. H. et al. CRISPR RNA-guided integrases for high-efficiency, multiplexed bacterial genome engineering. Nat. Biotechnol. 1–10 (2020) doi:10.1038/s41587-020-00745-y.

62. Rubin, B. E. et al. Species- and site-specific genome editing in complex bacterial communities. Nat. Microbiol. 7, 34–47 (2022).

63. Feiss, M. & Catalano, C. E. Bacteriophage Lambda Terminase and the Mechanism of Viral DNA Packaging. Madame Curie Bioscience Database [Internet] (Landes Bioscience, 2013).

64. Jiang, Y. et al. Multigene Editing in the Escherichia coli Genome via the CRISPR-Cas9 System. Appl. Environ. Microbiol. 81, 2506–2514 (2015).

65. Jiang, W., Bikard, D., Cox, D., Zhang, F. & Marraffini, L. A. RNA-guided editing of bacterial genomes using CRISPR-Cas systems. Nat Biotechnol 31, 233–239 (2013).

66. Meyer, A. J., Segall-Shapiro, T. H., Glassey, E., Zhang, J. & Voigt, C. A. Escherichia coli “Marionette” strains with 12 highly optimized small-molecule sensors. Nat. Chem. Biol. 15, 196–204 (2019).

67. Gibson, D. G. et al. Enzymatic assembly of DNA molecules up to several hundred kilobases. Nat Methods 6, 343–5 (2009).

68. Salis, H. M., Mirsky, E. A. & Voigt, C. A. Automated design of synthetic ribosome binding sites to control protein expression. Nat. Biotechnol. 27, 946–U112 (2009).

69. Croswell, A., Amir, E., Teggatz, P., Barman, M. & Salzman, N. H. Prolonged Impact of Antibiotics on Intestinal Microbial Ecology and Susceptibility to Enteric Salmonella Infection. Infect. Immun. 77, 2741–2753 (2009).

70. You, Y., Moreira, B. G., Behlke, M. A. & Owczarzy, R. Design of LNA probes that improve mismatch discrimination. Nucleic Acids Res. 34, e60 (2006).

