## Supplementary Information for "*In situ* targeted mutagenesis of gut bacteria"

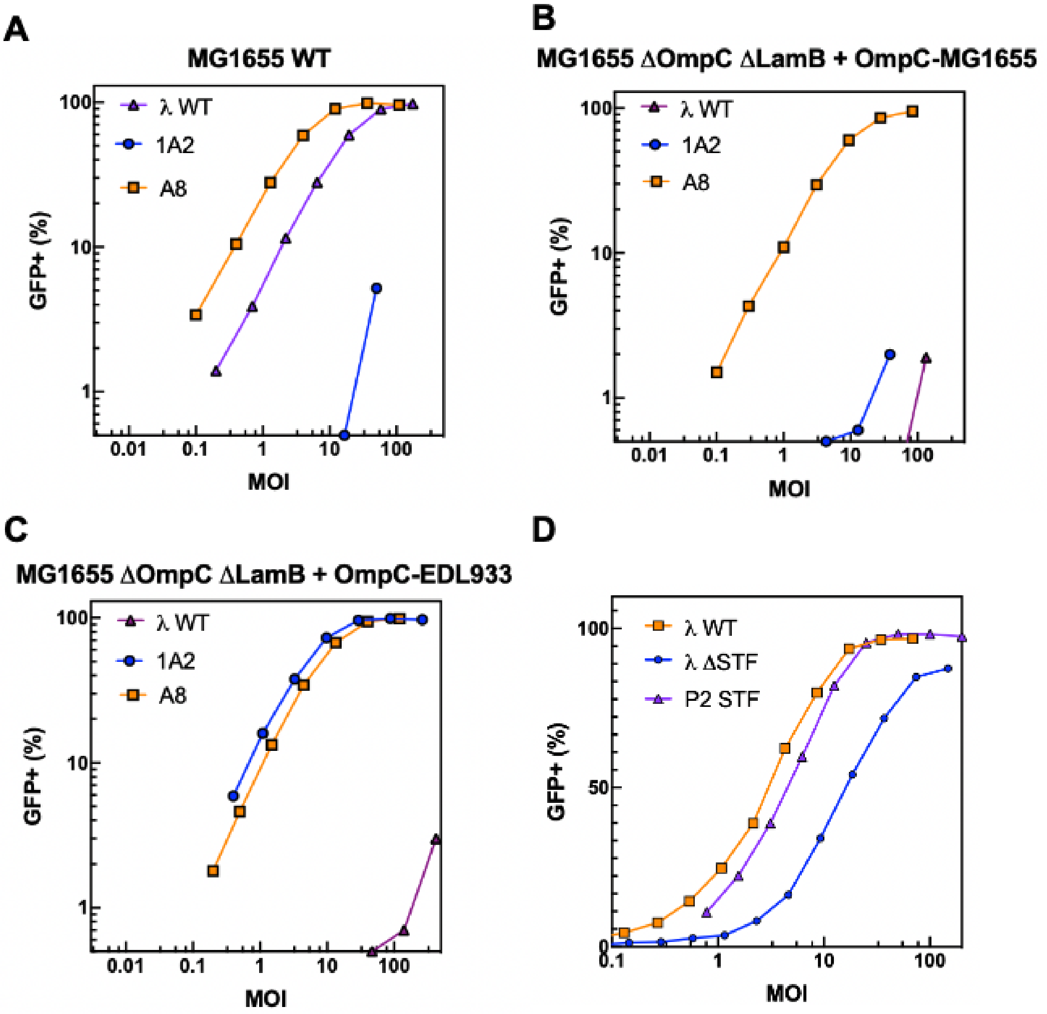


**Figure S1: Cell receptor analysis for efficient cosmid delivery.** Delivery efficiency assays of cosmids harboring Ur-λ wild-type (WT) gpJ and stf, or harboring gpJ chimeras and P2 STF. Transduction of a payload encoding a *sfGFP* fluorescent protein gene was measured in a flow cytometer (excitation: 488 nm, emission: 530/30 BP; Attune NxT Thermo Scientific) at different MOIs. **A)** Delivery efficiency into MG1655 WT, carrying the endogenous OmpC variant in the genome. **B)** and **C)** Delivery efficiency into a MG1655 strain deleted for both *lamB* and *ompC* genes, complemented with two different *ompC* variants on a plasmid (p1471 or p1472). **D)** Delivery efficiencies in *E. coli* MG1655 of λ particles carrying the WT λ STF (orange line), no STF (blue line) and λ-P2 STF chimera (p938) (purple line).

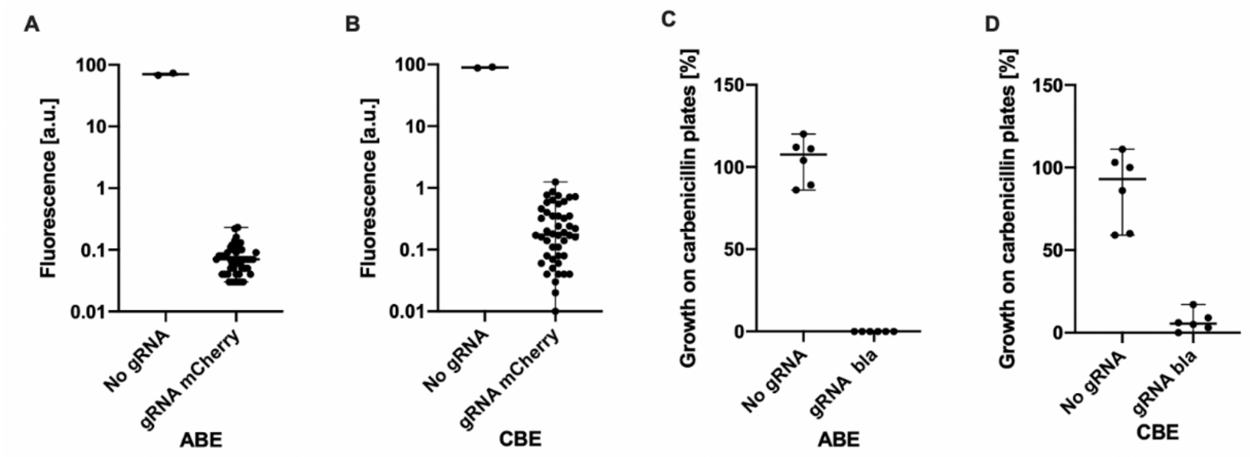


**Figure S2: Targeted base editing of the *E. coli* genome *in vitro* after DNA payload transformation and antibiotic selection. A)** Adenine base editing (ABE = ABE8e, plasmid p2325) of MG1655-*mCherry*. The experiment was performed in the presence (48 colonies analyzed) or absence (2 colonies analyzed) of a guide RNA targeting the active site of mCherry (tripeptide: M71, Y72, G73). **B)** Cytosine base editing (CBE = evoAPOBEC1-nCas9-UGI, plasmid p2326) of mCherry on MG1655-*mCherry* genome. The experiment was performed in presence (48 colonies analyzed) or absence (2 colonies analyzed) of a guide RNA inserting a stop codon at position Q114* into mCherry. Fluorescence of individual colonies was measured by flow cytometry (excitation: 561 nm, emission: 620/15 BP; Attune NxT Thermo Scientific). Dots represent single colonies after overnight incubation on chloramphenicol plates. **C)** Adenine base editing of β-lactamase (*bla*) in MG1655-*bla in vitro* (plasmid p1396). The experiment was performed in presence or absence of a guide RNA targeting the active site of β-lactamase (K73E or K73R). **D)** Cytosine base editing of β-lactamase in MG1655-*bla in vitro* (plasmid p2327). The CBE inserts a premature stop codon (Q37*) into the target gene, resulting in the re-sensitization of the bacterial population to β-lactam carbenicillin. Adenine and cytosine base editing of β-lactamase was analyzed by colony counting after overnight incubation on chloramphenicol (CA) and CA/carbenicillin agar plates at 30°C. The percentage growth was obtained by dividing the number of colonies on CA/carbenicillin plates by the number on CA plates. One dot represents one transformation after overnight incubation on plates.

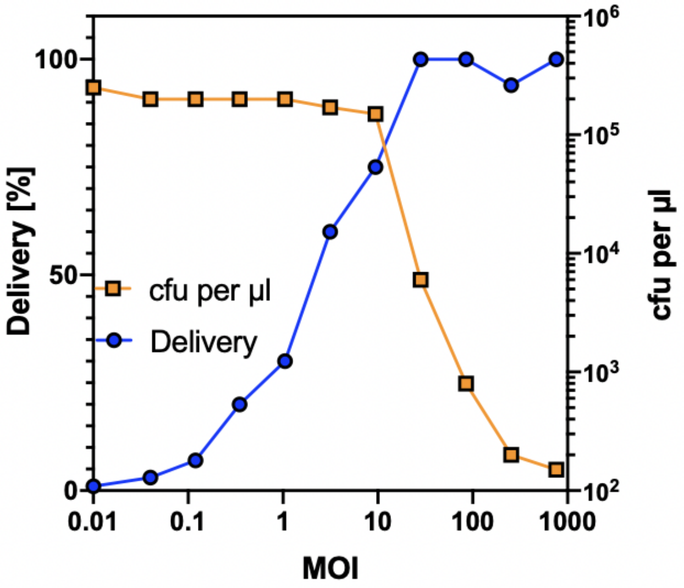


**Figure S3: Multiplicity of injection (MOI)-dependent delivery efficiency and corresponding adenine base editing (ABE).** Delivery efficiency was obtained by colony counting after overnight incubation on LB and LB (chloramphenicol) agar plates. Left y axis: The percentage delivery was obtained by dividing the number of colonies on chloramphenicol plates by the number on LB plates. Right y axis: Base editing of β-lactamase in strain MG1655-*bla* diminishes cell growth on carbenicillin plates (colony-forming units cfu per µl).

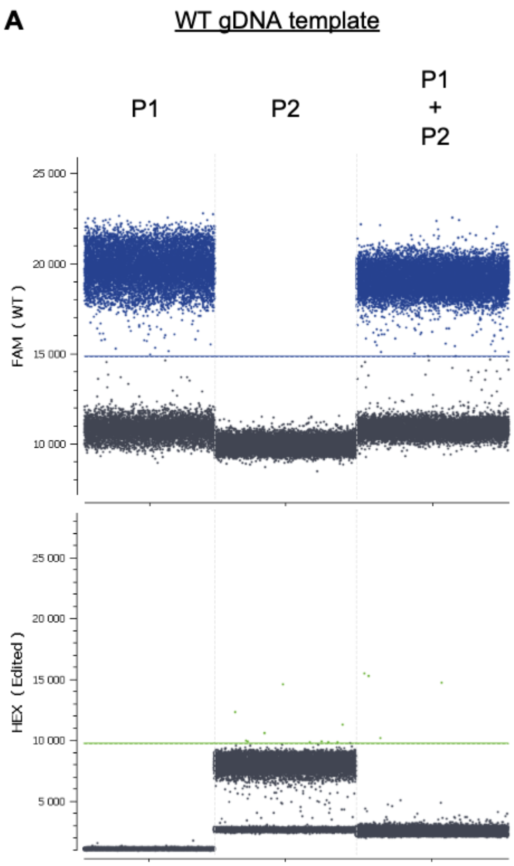


**Figure S4: Validation of ddPCR Taqman assay.** Probes P1 (wild-type WT target) and P2 (base-edited target) were assessed for their specificity, either individually or when mixed together at a 1:1 molar ratio, using purified genomic DNA (gDNA) from wild-type WT (**panel A**) and *in vitro*-edited (**panel B**) MG1655-*bla* as template.

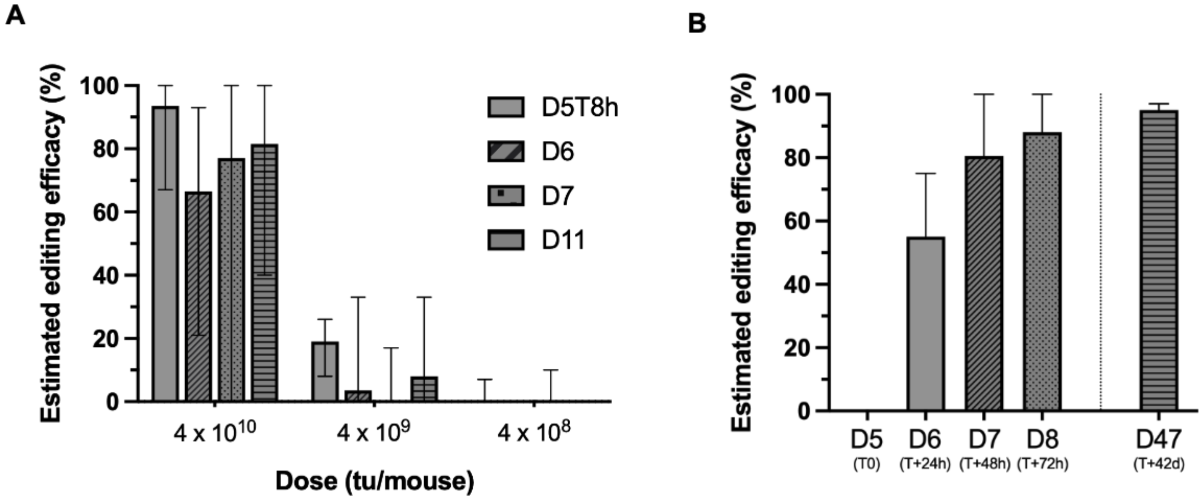


**Figure S5: Estimated efficacy of targeted adenine base editing on the *E. coli* genome in gut of BALB/c mice after cosmid treatment using a non-replicative payload.** Estimation of the editing efficacy as measured by repatching individual colonies onto agar plate with or without carbenicillin, for the initial dose-response experiment (**panel A**), and for the experiment looking at cumulative effects (**panel B**). Bars represent the group median, with 95% confidence interval.

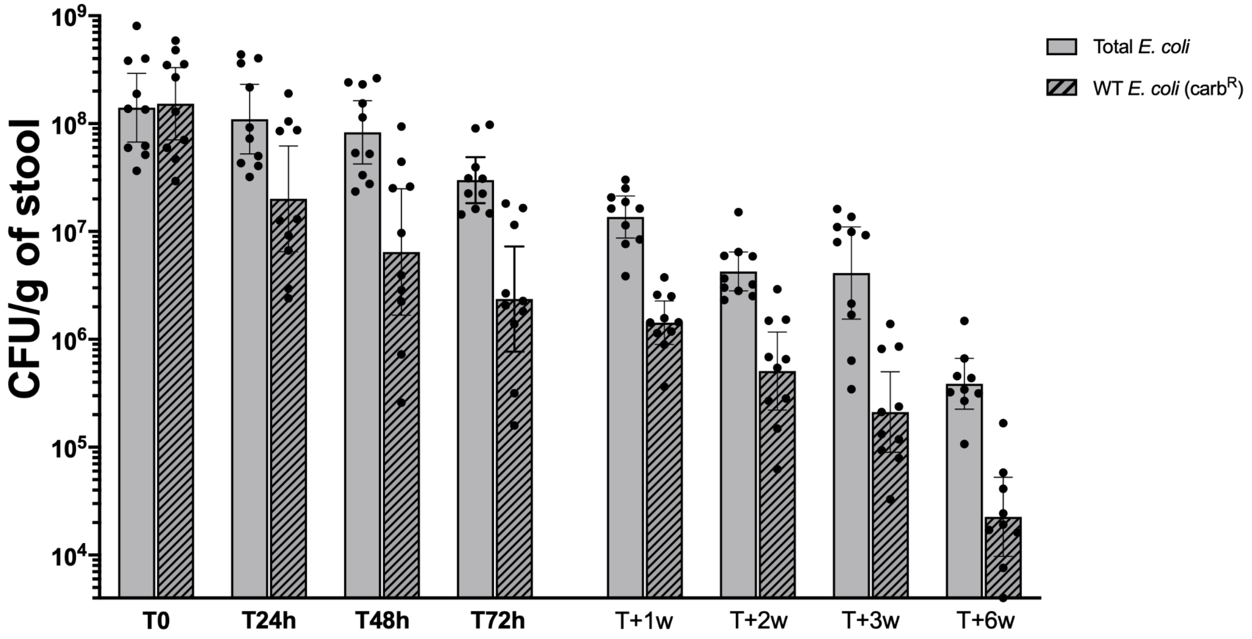


**Figure S6: *E. coli* s21052 colonization levels over time in the streptomycin-treated BALB/c model during treatment with base-editing cosmid particles.** Total *E. coli* bacteria were quantified by plating resuspended stool samples into Drigalski plates supplemented with streptomycin. Wild-type WT (ie, non-edited) *E. coli* s21052 were quantified by plating samples onto Drigalski plates with carbenicillin.

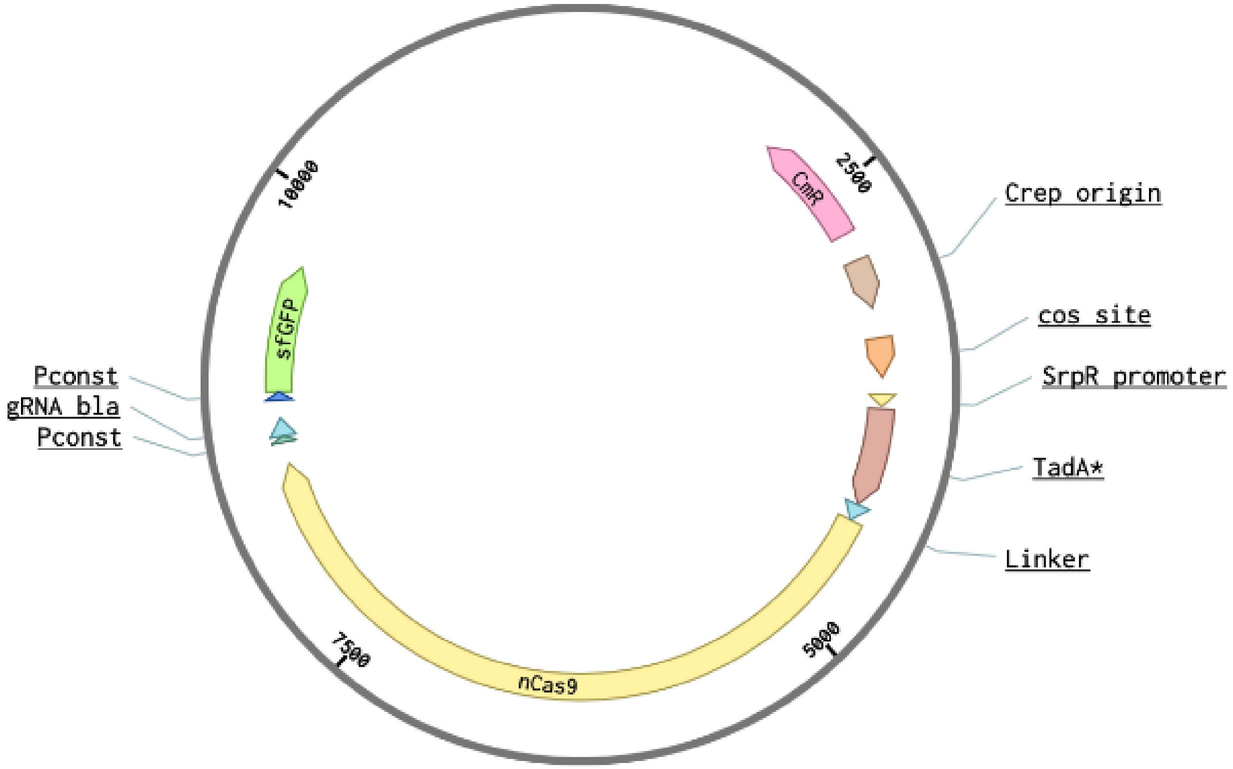


**Figure S7: Plasmid map of the non-replicative cosmid encoding the adenine base editor and a guide RNA targeting the active site of β-lactamase (*bla*).** Packaging into λ phage particles is enabled by the cohesive end site (*cos)* of the λ genome. The adenine base editor is a fusion gene of *TadA** and *nCas9* under the SrpR promoter. The guide RNA targeting the *bla* gene is under a constitutive promoter. The plasmid carries a constitutively-expressed chloramphenicol resistance gene (CmR) and the non-replicative primase origin of replication (Crep). In addition, the plasmid carries the *sfGFP* gene under a constitutive promoter to investigate cosmid delivery efficiencies. The total plasmid size is 10,747 bp.

**Table S1: Genotypes of *E. coli* strains used in this study.**

| Strain | Genotype | Supplier |
|---|---|---|
| DH10B | *F– mcrA Δ(mrr-hsdRMS-mcrBC) φ80lacZΔM15 ΔlacX74 recA1 endA1 araD139 Δ (ara-leu)7697 galU galK λ– rpsL(StrR) nupG* | Thermo Scientific |
| CY2120 | *BW25113 Δ9 λcI857 PaPa Sam7 Δcos lamB-* | University of Illinois |
| CY-1A2 | *BW25113 Δ9 λcI857 Sam7 Δcos gpJ-1A2 ∆stf lacZ::SrpR lamB-* | This study |
| CY-A8 | *BW25113 Δ9 λcI857 Sam7 Δcos gpJ-A8 ∆stf lacZ::SrpR lamB- ∆OmpC* | This study |
| CY-Ur-λ | *BW25113 Δ9 λcI857 orf401::orf314 Sam7 Δcos lamB-* | This study |
| MG1655 | *K-12 F- λ- ilvG- rfb-50 rph-1* | Pasteur Institute |
| MG1655-*ΔLamB* | *K-12 F- λ- ilvG- rfb-50 rph-1 ΔLamB* | This study |
| MG1655-*ΔLamB*-ΔompC | *K-12 F- λ- ilvG- rfb-50 rph-1 ΔLamB ΔompC* | This study |
| MG1655-*mCherry* | *K-12 F- λ- ilvG- rfb-50 rph-1 mCherry* | Pasteur Institute |
| MG1655-*bla* | *K-12 F- λ- ilvG- rfb-50 rph-1 rpsLK42R wbbL::bla-rfp* | This study |
| s14269 | *K-12 F- λ- ilvG- rfb-50 rph-1 rpsLK42R ompC-EDL933* | This study |
| s14269-bla | *K-12 F- λ- ilvG- rfb-50 rph-1 rpsLK42R wbbL::bla-rfp ompC-EDL933* | This study |
| s21052 | *K-12 F- λ- ilvG- rfb-50 rph-1 rpsLK42R wbbL::bla-rfp ompC-EDL933 rpslK42R* | This study |
| *E. coli* CFT073 | *O6:K2:H1* | Pasteur Institute |

**Table S2: List of plasmids used in this study.**

| Name | Plasmid | Class | Resistance | Source | Notes |
|---|---|---|---|---|---|
| p2325 | p15A-cos-pSrpR-RBS-ABE8e gRNA1(mCherry) | Cosmid | CamR | This study | Encodes ABE targeting mCherry in a constitutive origin of replication |
| p2326 | p15A-cos-pSrpR-RBS-CBE gRNA2(mCherry) | Cosmid | CamR | This study | Encodes CBE targeting mCherry in a constitutive origin of replication |
| p1396 | p15A-cos-pSrpR-RBS-ABE8e gRNA4(*bla*) | Cosmid | CamR | This study | Encodes ABE targeting bla in a constitutive origin of replication |
| p2327 | p15A-cos-pSrpR-RBS-CBE gRNA5(*bla*) | Cosmid | CamR | This study | Encodes CBE targeting bla in a constitutive origin of replication |
| p2328 | pCrep-cos-pSrpR-RBS-ABE8e gRNA4(*bla*) | Cosmid | CamR | This study | Encodes ABE targeting bla in a conditional origin of replication |
| p1324 | pCrep-cos-sfGFP | Cosmid | CamR | This study | Encodes sfGFP in a conditional origin of replication |
| p513 | p15A-cos-sfGFP | Cosmid | CamR | This study | Encodes sfGFP in a constitutive origin of replication |
| p938 | pSC101-pPhlF-RBS-stf P2 | Stf | KanR | This study | Encodes an inducible λ-P2 STF chimera |
| p1471 | pEco-OmpC MG1655 G1 | Receptor | KanR | This study | Encodes OmpC from MG1655 |
| p1472 | pEco-OmpC G17 | Receptor | KanR | This study | Encodes OmpC from EDL933 |
| p2076 | pIncW-RARE7-RBS-primase | Crep system | TpR | This study | Encodes a constitutive primase gene |
| p1321 | pSC101-pPhlF-RBS-primase | Crep system | KanR | This study | Encodes an inducible primase gene |

**Table S3: List of gRNAs used in this study.**

| Name | Editor | gRNA | Position | Target |
|---|---|---|---|---|
| gRNA1 | ABE | CGTACATAAATTGCGGGCTC | 4A, 6A, 8A | mCherry (M71T, Y72H) |
| gRNA2 | CBE | CACTCAGGACTCCTCCCTGC | 5C | mCherry (Q114*) |
| gRNA3 | CBE | TAGAACCGTACATAAATTGC | 6C, 7C | mCherry (G73N or G73D) |
| gRNA4 | ABE | ACTTTTAAAGTTCTGCTATG | 1A, 7A, 8A | β-lactamase (K71E or K71R) |
| gRNA5 | CBE | GATCAGTTGGGAGCCCGTGT | 4C | β-lactamase (Q37*) |

**Table S4: Selection of oligonucleotides used for plasmid sequencing and ddPCR.** Probes P1 and P2 contained a different fluorophore (FAM or HEX), as well as carefully positioned Locked Nucleic Acid bases (LNA; symbolized by the base A, T, C, or G preceded by a "+" sign in the sequences above).

| Name | Oligonucleotide sequence | Gene |
|---|---|---|
| F1 | ATGGTTTCCAAGGGCGAGG | mCherry |
| R1 | TTATTTGTACAGCTCATCCATGCC | mCherry |
| F2 | ATGAGTATTCAACATTTCCGTGTCGC | *Bla* |
| R2 | TTACCAATGCTTAATCAGTGATGC | *Bla* |
| F3 | GGATCTCAACAGCGGTAAG | *Bla* (ddPCR, primer) |
| R3 | GGCATCAACACGGGATAATA | *Bla* (ddPCR, primer) |
| P1 | FAM-CT+TT+T+A+AA+GTT+C+T+GC | *Bla* (ddPCR, probe 1) |
| P2 | HEX-CT+TT+T+G+AAGTT+CT+GC | *Bla* (ddPCR, probe 2) |
